# Integrated Spatial Multi-omic Profiling Identifies HSV-associated Inflammatory Macrophage Niches Linked to Oncolytic Virotherapy Response in Melanoma

**DOI:** 10.64898/2026.05.20.726697

**Authors:** Emma Wagner, Samuel Legg, Christopher J Applebee, Julian Padget, Banafshé Larijani, Amanda R Kirane

**Affiliations:** Division of General Surgery, Department of Surgery, Section of Surgical Oncology, Stanford University School of Medicine, Stanford, CA, USA; Department of Computer Science, University of Bath, Bath, UK; Cell Biophysics Laboratory, Department of Life Sciences, Consortium for Precision Health, University of Bath, Bath, UK; Department of Computer Science, Consortium for Precision Health, University of Bath, Bath, UK

**Keywords:** Cancer immunotherapy, melanoma, oncolytic virus, T-VEC, immune checkpoint interaction, functional proteomics, digital spatial profiling

## Abstract

**Background:** Primary and secondary resistance to immune checkpoint blockade (ICB) remains a critical challenge in advanced melanoma. Oncolytic Viruses (OV) selectively lyse tumor cells while generating systemic anti-tumor immune responses with minimal side effects. Yet their clinical use is limited to refractory melanoma patients and are only given in combination with second-line ICB regimens. ICB can both help and hinder OV efficacy depending on the source of checkpoint interactions across the tumor-immune microenvironment (TiME). However, functional checkpoint interactions are typically inferred from gene or protein expression and rarely contextualized within myeloid- and antigen presenting cell-associated immune niches during OV therapy, despite these populations dominating melanoma TiMEs and serving as key regulators of anti-viral immunity.

**Methods:** An integrated multi-omics framework combining Nanostring GeoMx digital spatial profiling (DSP), COMET sequential immunofluorescence (seqIF) and functional oncology mapping (FuncOmap) was applied to melanoma patient tissues collected pre- and post-neoadjuvant Talimogene Laherparepvec (T-VEC) to characterize immune remodeling and directly quantify checkpoint interaction dynamics associated with pathologic responses to OV therapy.

**Results:** T-VEC induced broad lymphocyte- and myeloid-associated immune transcriptional activation across melanoma TiMEs; however, pathologic responses could not be defined by bulk transcriptomics or cellular deconvolution alone. COMET seqIF analysis identified that HSV-associated M1/APC-like tumor-associated macrophages (TAMs) were enriched in complete pathologic response (CR) tissues and were a major source of PD-1/PD-L1 interaction niches. While partial (PR) and non-pathologic response (NR) tissues retained melanoma-centered PD-1/PD-L1 interaction niches and were enriched for B cell and M2-like TAM populations. FuncOmap analysis indicated that post-T-VEC PD-1/PD-L1 interaction states were consistently elevated in tumor bed, but not in lymph node tissues, across all pathologic response groups. Suggesting that immune checkpoint interactions may benefit T-VEC therapeutic responses depending on their spatial and immune context relative to OV infection.

**Conclusions:** These findings highlight the importance of integrated transcriptomic and functional proteomic analyses for resolving the spatial distribution and functional status of immune niches during OV therapy. Resolving PD-1/PD-L1 interaction states to specific M1/APC-like TAM and B cell niches may define mechanisms of responses and resistance to OV therapy.

## Introduction

Immune checkpoint blockade (ICB) has revolutionized overall responses rates in melanomas; however, a significant number of patients still acquire resistance or show primary resistance to ICBs^1–4^. Patients with refractory melanomas are often recommended to pursue second-line or combination ICB regimens or tumor-infiltrating lymphocyte (TIL) therapy which carry high risk of severe immune-related adverse events (irAEs) and toxicities^5,6^. Oncolytic viruses (OV) are promising alternative systemic immunotherapies to ICB and TIL treatment due to their ability to selectively lyse cancer cells whilst sparing host tissues and thereby minimizing toxicities and irAEs^7–11^. Talimogene laherparepvec (T-VEC) is a genetically modified herpes simplex virus type 1 (HSV-1) OV strain that was approved for treatment of advanced melanoma patients in 2015 following the results of the OPTiM phase III randomized trial ^12–17^. However, in the decade since its approval, real-world T-VEC clinical use in melanomas is often restricted to patients who have failed first-line ICB and T-VEC is only given in combination with second-line ICB therapies^18–20^.

Multiple clinical trials investigating the benefit of T-VEC as a monotherapy or combination with ICB in treatment naïve and heavily pretreated advanced melanomas have had mixed results. Where T-VEC as a monotherapy or combination with ICB improved responses rates in advanced refractory melanomas yet did not significantly improve overall survival in treatment-naïve advanced melanomas^21^. ICB therapeutic efficacy has long been attributed to increased infiltration and prolonged activity of T-lymphocytes, particularly CD8^+^ cytotoxic T lymphocytes, as the key drivers of anti-tumor immunity in melanoma tumor-immune microenvironments (TiME). However, durable responses to OV therapy are dependent on concurrent activation of anti-tumor and anti-viral immunity and the benefit of immune checkpoint inhibition in anti-viral immunity is less clear^22^. Functional interactions between programmed death receptor 1 (PD-1) and its ligand (PD-L1) can both help and hinder anti-viral immunity in response to OV infection depending on the interaction sources in the TiME^23,24^. Melanoma TiMEs include heterogenous mixtures of lymphocyte and antigen presenting cell (APC) populations, where tumor-associated macrophages (TAMs) which are the most abundant type of APCs and are also key regulators of immune responses to HSV infection^25–27^.

Recent advances in digital spatial profiling (DSP) using transcriptomic and multiplex imaging platforms now enable integrated characterization of TiMEs at the genomic and proteomic level. These spatial analyses have accelerated discovery of multicellular niches whose functional significance to driving immunotherapy responses depends on contextualizing the spatial organization of the surrounding melanoma TiMEs. However, when it comes to defining the contribution of these immune niches to PD-1/PD-L1 interaction states, these spatial multi-omics platforms traditionally infer PD-1/PD-L1 interactions from their relative gene or protein expression in the TiME which is not indicative of their functional status^28^. Moreover, the functional interaction states of immune cells relative to HSV infection during T-VEC therapy remain poorly understood.

We present here, for the first time, an integrated DSP approach combining Nanostring GeoMx transcriptomics, COMET sequential immunofluorescence (seqIF) and functional oncology mapping (FuncOmap) of checkpoint interactions to characterize immune remodeling and directly quantify PD-1/PD-L1 interaction dynamics within the melanoma TiME in response to T-VEC therapy. Applying this multimodal framework, we identified T-VEC therapy induces broad lymphocyte- and myeloid-associated immune activation although pathologic complete response (CR) was associated with enriched HSV-associated M1 TAM populations linked to localized PD-1/PD-L1 niches. Whereas partial (PR) and non-pathologic responses (NR) to T-VEC retained melanoma-centered PD-1/PD-L1 niches and were enriched with B cell and M2-like TAM populations. Altogether, this multimodal spatial framework provides functional insight into the melanoma TiME features potentially associated with response and resistance to OV therapy.

## Methods

### Patient Samples and Study Design

Patient samples used in this study were collected from six patients with stage II melanoma before and after treatment with neoadjuvant T-VEC therapy as previously described^29^. Briefly, patient samples were collected at baseline and one week post final T-VEC injection. Baseline samples were collected as punch biopsies up to 30 days before receiving the first T-VEC dose. T-VEC dosing was dependent on patient lesion size, all patients were given a loading dose (10^6^ PFU/mL) 3 weeks before beginning treatment doses (10^8^ PFU/mL). T-VEC treatment doses were administered on days 1, 21, 35 and 49 for all patients. Post-T-VEC samples were procured via wide local excision and sentinel lymph node (LN) biopsy on week post final treatment dose. Patient tissues were preserved as formalin-fixed paraffin-fixed (FFPE) samples, 2mm cores were obtained from tumor centers and skin-tumor periphery in pre-T-VEC and from central tumor beds and SLNs for post-T-VEC samples were used to construct a tissue microarray (TMA) FFPE block composed of 24 tissue cores (x4 tissue cores per patient). Where each patient was represented by two pre-T-VEC tissue cores (x1 central tumor (CT) and x1 skin-tumor interface (STI)) as well as two post-T-VEC cores (x1 central tumor bed (CT) and x1 SLN). Five sequential TMA slides were cut per patient (4µm thickness) for downstream hematoxylin and eosin (H&E) staining, Nanostring GeoMx DSP, FuncOmap and COMET seq analysis. H&E-stained sections were used to confirm CT (pre- and post-T-VEC, STI (pre-T-VEC) and LN (post-T-VEC) regions and for scoring pathologic response and were reviewed by three independent dermatopathologists.

### Nanostring GeoMx DSP Transcriptomic Profiling

The FFPE TMA slide was processed, and antigen retrieval was performed per Nanostring GeoMx DSP guidelines^30^. Manual region of interest (ROI) selection was guided by CD3, CD68, DAPI and aSMA fluorescent antibody staining to select immune enriched verses stromal regions across tissue cores per GeoMx guidelines. A total of 56 ROIs were selected across 24 tissue cores (∼3-4 ROIs per core) and were collected for bulk transcriptomics using the whole human transcriptome atlas (WTA) per GeoMx DSP guidelines. Raw count data was imported into R (v4.5.2) and converted into a SpatialExperiment object for analysis using the standR framework^31^. Quality control (QC) steps were performed at the gene and ROI level. Lowly expressed genes were removed using a logCPM-based filtering threshold, excluding genes with minimal expression across 90% of ROIs. ROIs containing fewer than 100 nuclei were excluded following inspection of QC steps. Negative control probes were used for technical QC but excluded from downstream expression analyses. Gene expression data were normalized using trimmed mean of M-values (TMM) normalization followed by voom transformation to generate log2-counts per million (logCPM) values for downstream analyses.

### Differential Expression and Pathway Analysis

Differential expression (DE) analyses were performed using the limma-voom framework. Gene-level counts were converted into DGEList objects, filtered using filterByExpr, and normalized using TMM normalization. Linear modeling was performed following voom transformation to account for mean-variance relationship in sequencing count data. To account for repeated sampling of multiple ROIs per patient, patient identity was incorporated as a blocking factor using duplicateCorrelation. Three primary contrasts were evaluated; i) CT pre-vs post-T-VEC treatment, ii) LN verses CT post-T-VEC, and iii) STI verses CT pre-T-VEC tissues. Moderated t-statistics were computed using empirical Bayes shrinkage (eBayes). DE genes were defined using a false discovery rate (FDR) threshold <0.05, with additional log2 fold-change thresholds applied for visualization where appropriate. Gene set enrichment analysis (GSEA) was performed using curated immune-related gene sets from the molecular signatures database (MSigDB), including Hallmark (H) and immune-focused C7 collections^32,33^. Gene sets were restricted to genes retained following expression filtering to maintain consistency between gene-level and pathway-level analyses. Enrichment significance was assessed using the fry function from limma, and representative immune-associated pathways were selected for downstream visualization and interpretation.

### Spatial Cellular Deconvolution

Spatial cellular deconvolution was performed using the SpatialDecon framework and safeTME reference matrix to estimate immune cell composition within individual ROIs^34^. Normalized expression data were prepared for deconvolution according to the SpatialDecon workflow, and estimated cell-type proportions were calculated per ROI. Differences in cell type proportions were transformed using arcsine square-root transformation, and linear modelling was performed with patient identity included as a blocking factor. Statistical inference was determined using false discovery rate (FDR)-adjusted p values.

### COMET seqIF Imaging

FFPE patient tissue microarray was de-waxed and antigen retrieval was performed by placing the slides in Dewax and HIER Buffer H (Epredia, catalogue number: TA-999-DHBH) inside the PT Module (Epredia, catalogue number: A80400011) for 20 mins at 95°C. The processed TMA slide was immediately placed into Multi-staining Buffer (Lunaphore Technologies SA, catalogue number: BU06) until use. The TMA slide was transferred to the COMET instrument where a microfluidic COMET Chip (Lunaphore Technologies SA, catalogue number: MK03) was sealed on top of the processed TMA tissues by the COMET instrument to form a closed reaction chamber. FFPE tissue sections were stained by reagents delivered via the microfluidic channels according to the pre-loaded seqIF protocols. Automated seqIF protocols were generated using the COMET Control software and consisted of iterative autofluorescence, staining, imaging and elution cycles^35^. Primary antibodies, secondary antibodies and nuclear counterstain Hoechst 33342 were diluted to their desired concentrations using multi-staining buffer (MSB, Lunaphore Technologies SA, catalogue number: BU06). Slides were imaged at 20X magnification within a 12.5 x 12.5mm imaging area by the integrated fluorescent microscope using DAPI (exposure time: 25 ms), TRITC (exposure time: 250 ms) and Cy5 (exposure time: 400 ms) channels for every imaging cycle. Primary and secondary antibodies were incubated for 8 and 16 minutes respectively in each cycle based on prior optimization. After each imaging cycle, antibodies were eluted using the default settings for each elution cycle. Once completed, the COMET seqIF protocol generated a multi-layer OME-TIFF consisting of the stitched and aligned images from each imaging cycle.

### COMET seqIF Image Analysis

COMET seqIF OME-TIFF was opened in Horizon™ Viewer software (Lunaphore Technologies SA). Background-subtracted OME-TIFFs were exported for downstream analysis in JupyterLab using the image processing pipeline SPACEc where individual TMA cores were extracted for downstream analyses^36^. Cell segmentation was performed independently for each core using Mesmer deep learning segmentation algorithm implemented within SPACEc. Nuclear segmentation was performed using Hoechst staining, while whole-cell boundaries were refined using broadly expressed membrane and cytoplasmic markers to capture immune, stromal and tumor compartments. Segmentation quality was assessed by visual inspection of representative image regions with overlaid cell boundaries. Per-core segmentation feature tables were imported into JupyterLab and concatenated using the python-based SPACEc preprocessing workflow. Low-quality segmented objects were removed based on cell area and nuclear Hoechst intensity thresholds. Marker intensities were normalized independently within each core using z-score transformation, and noisy cells were removed using marker intensity and z-count thresholding. Single cell analyses were performed using Scanpy^37^ and Spacec^36^ workflows. Dimensionality reduction was performed using uniform manifold approximation and projection (UMAP) and unsupervised clustering was performed using the Leiden algorithm. Broad immune and tumor populations were annotated according to canonical marker expression patterns, including melanoma cells, macrophages, T cells, B cells, stromal cells, and myeloid populations. Major immune lineages were subsequently subset and re-clustered to resolve finer phenotype annotations including M1/APC-like, M2-like and intermediate tumor associated macrophage (TAM) populations, as well as CD4+ T cell, CD8+ T cell and NK/T cell lymphocyte populations. Clusters lacking coherent lineage-associated marker expression were designated as unresolved and excluded from downstream analyses.

### HSV and PD-1/PD-L1 Annotations

Given T-VEC is derived from a herpesvirus (HSV) backbone, HSV-associated cell states were identified using combined expression of three non-redundant HSV proteins corresponding to early, intermediate and late stages of the HSV life cycle; ICP0 (intermediate-early stage), ICP8 (early stage) and HSV glycoprotein D (HSVgD, late stage). Concordant expression of all three proteins was determined using 99^th^ percentile fluorescence intensity threshold that was used to generate a composite score for HSV positivity that was used for downstream spatial analyses. PD-1/PD-L1-associated cell states were also defined using 95^th^ percentile-based fluorescence intensity thresholds applied independently within each tissue core. PD-1-high and PD-L1-high annotations were subsequently incorporated into downstream spatial and proximity analyses.

### Spatial Proximity and Neighborhood Analyses

Pairwise spatial proximity analyses were performed between PD-1-high source and PD-L1-high target cell populations across matched post-T-VEC CT and LN tissues. For each tissue core, nearest-neighbor distances were calculated between all PD-1-high source cells and the nearest PD-L1-high target cells using cell centroid coordinates. Analyses were restricted to cell populations containing at least 20 PD-1-high source cells and 20 PD-L1-high target cells per interaction pair to reduce variation from sparsely represented populations. For each source-target cell type combination, median nearest-neighbor distance, mean nearest-neighbor distance, and the proportion of source cells within a 5-pixel radius of a target population were calculated. Exploratory proximity screening was used to identify recurrent biologically relevant PD-1/PD-L1-associated cellular interactions involving melanoma cells, TAM subsets, NK/T-like cells and B cells for focused visualization and interpretation. Neighborhood composition analyses were subsequently performed to characterize the local immune and tumor microenvironment surrounding HSV-associated M1/APC-like TAM populations within post-T-VEC CT cores. HSV-high M1/APC-like TAMs were used as anchor cells, and neighboring cells within a 50-pixel radius were identified using radius-based nearest-neighbor analysis. Neighboring cells were grouped into broad biological categories, including TAM, CD8 T cell, NK/T cell, B cell, melanoma and stromal populations. Neighborhood compositions were converted to proportional abundances and averaged across anchor cells within each tissue core for comparative analyses across pathological response groups.

### FuncOmap Analysis

Two sequential FFPE TMA slides underwent antigen retrieval, tyramide signal amplification, and labelling with donor and acceptor antibodies and acquisition as previously described^38^. PD-1/PD-L1 interactive states are calculated as FRET-efficiency (Ef) and reflect the reduction of lifetime of donor (PD-1) in presence of the acceptor (PD-L1) chromophores. Per-pixel Ef maps are generated using FuncOmap software pipeline as previously described, where each pixel represents the calculated Ef value corresponding to PD-1/PD-L1 interaction. Distribution of all Ef values across each TMA tissue core are quantified using violin plots, corresponding to approximately 2.6×10^5^ datapoints per (512×512 pixels) FuncOmap region. The global violin plots of each 1mm core highlights the distribution of all 5.2×10^6^ Ef data points captured per tissue core. Given that Ef distributions are non-Gaussian, the upper quartile (Q3) of the global Ef distribution and its population density (0-1) is used to represent regions with the highest PD-1/PD-L1 interaction states across tissue cores^38,39^.

### Statistical Analyses

All statistical analyses were performed in R (v4.5.2) or Python (JupyterLab). Differential expression and pathway enrichment analyses were performed using the limma framework with Benjamini–Hochberg correction to control the false discovery rate. Cellular proportion analyses were performed using the propeller framework with FDR adjustment. Correlation analyses were performed using Spearman’s rank correlation. All statistical tests were two-sided, and statistical significance was defined as FDR <0.05 unless otherwise specified. Given the limited cohort size, analyses were considered exploratory and hypothesis-generating.

## Results

### T-VEC induces transcriptional immune remodeling across tumor and lymph node compartments

Spatial bulk transcriptomics was performed using Nanostring GeoMx DSP to characterize the tumor-immune microenvironment (TiME) across central tumor (CT), skin-tumor interface (STI) and lymph node (LN) tissues pre- and post-T-VEC therapy. Differential gene expression analysis comparing pre- and post-T-VEC central tumor (CT) regions identified widespread transcriptional changes including upregulation of several immunoglobulin genes including IGKC and IGHG1-4 in post-T-VEC samples (Figure 1A). Principal component analysis demonstrated separation of this immunoglobulin signal with post-T-VEC CT regions. Additional characterization of the differentially expressed genes was performed using gene set enrichment analysis (GSEA) using curated immune-related signatures. Post-T-VEC CT regions demonstrated enrichment of CD8^+^ T cell, B cell and macrophage activation-associated signatures compared with pre-T-VEC CT tissue (Figure 1B).

**Figure 1.**
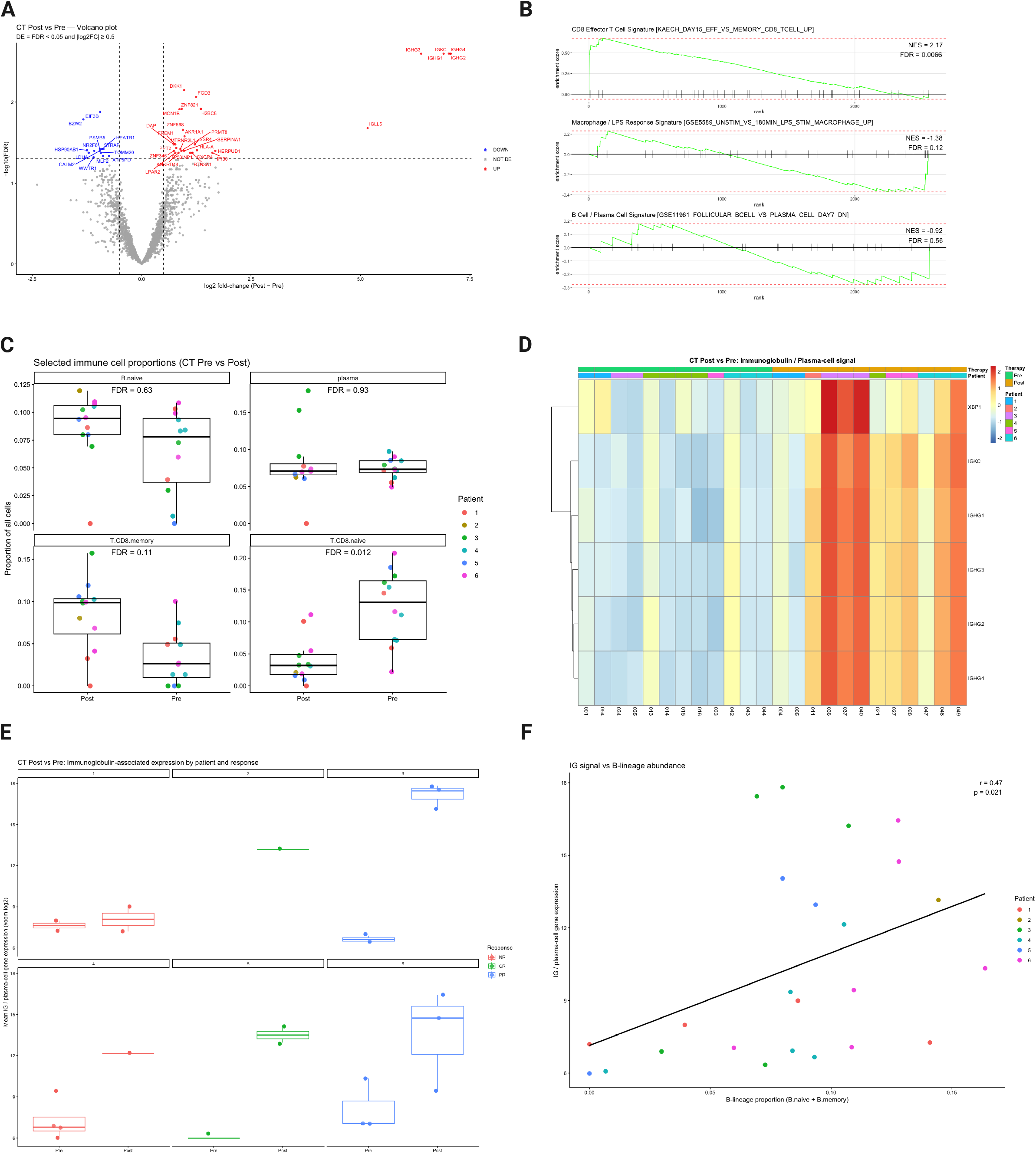
Spatial transcriptomic profiling identifies immune remodeling within central tumor regions following T-VEC therapy. A) Volcano plot showing differential gene expression between post-treatment and pre-treatment central tumor (CT) regions following T-VEC therapy. Selected significantly upregulated immunoglobulin associated genes are highlighted. B) Gene set enrichment analysis (GSEA) of post-treatment versus pre-treatment CT regions demonstrating enrichment of immune-associated transcriptional programs, including cytotoxic T-cell, macrophage, and B/plasma-cell signatures. C) Spatial deconvolution analysis showing relative immune cell-type proportions across pre-treatment and post-treatment CT regions. D) Heatmap of selected immunoglobulin-associated transcripts across CT regions demonstrating coordinated enrichment in post-treatment tissues. E) Per-patient summary of immunoglobulin-associated expression scores across CT regions before and after therapy. F) Correlation between immunoglobulin-associated expression scores and estimated B-lineage cell proportions across CT regions.

Spatial cellular deconvolution analyses were subsequently performed to identify TiME cell populations associated with these transcriptional changes using the safeTME reference matrix. These analyses identified elevated B naïve cell and CD8^+^ T memory cell populations in post-compared to pre-T-VEC CT regions, although these changes were not significant. No significant changes in the proportion of plasma cells from pre-to post-T-VEC were detected (Figure 1C). The proportion of CD8^+^ T naive cells was significantly (p=0.012) decreased in post-T-VEC CT regions compared to pre-T-VEC CT regions. Patient-level analyses confirmed the expression immunoglobulin (Ig)-associated genes were elevated in post-T-VEC compared to pre-T-VEC CT regions across all six patients. However, the highest increases in Ig-associated genes were seen in CT regions collected from patients with partial pathologic responses (PR) (Figure 1D, E). Increases in Ig/plasma cell-associated gene expression positively correlated with the proportion of combined (naive + memory) B cell populations across CT regions (r=0.47, p=0.021) (Figure 1F).

Differential gene expression analyses were also performed between pre-T-VEC CT and STI samples as well as post-T-VEC CT and LN samples to investigate regional differences in immune organization and transcriptional activity. Significant differences in gene expression were not observed between pre-T-VEC CT and STI tissues, however, LN tissues showed significant upregulation of multiple lymphocyte-associated, and antigen presentation-related genes compared to post-T-VEC CT regions (Figure 2A). GSEA revealed CD8^+^ T effector cell and macrophage-associated signatures were depleted in LN compared to post-T-VEC CT tissues, while CD8^+^ T naïve/memory cell and B memory cell signatures were enriched in LN verses post-T-VEC CT tissues (Figure 2B). Spatial cellular deconvolution of these tissue regions did not identify significant differences in the proportion of B naïve cell or macrophage populations, however, the proportions of CD4^+^ T naïve (p=0.031) and CD4^+^ T memory (p=0.042) cell populations were significantly increased post-T-VEC in LN compared to CT tissues (Figure 2C). Per-patient level analysis of DE genes confirmed that broad transcriptional activation of T lymphocyte-associated genes in LN compared to post-T-VEC CT tissues were consistent across all six patients (Figure 2D).

**Figure 2.**
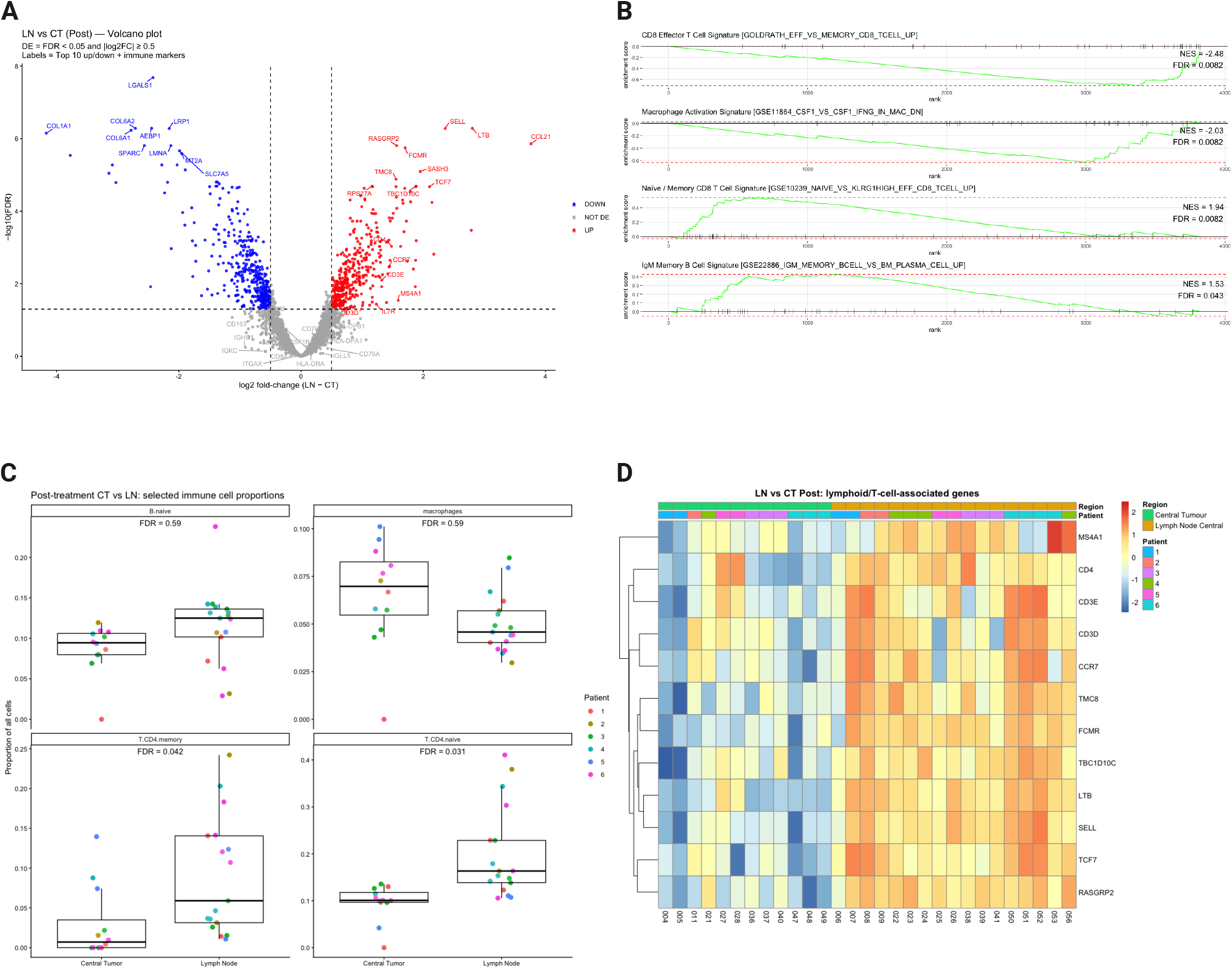
Lymph node and tumor regions exhibit distinct immune transcriptional programs following T-VEC therapy. A) Volcano plot showing differential gene expression between post-treatment lymph node (LN) and post-treatment central tumor (CT) regions. B) Gene set enrichment analysis demonstrating enrichment of lymphocyte-, macrophage-, and antigen presentation-associated transcriptional programs within LN tissues relative to CT regions. C) Spatial deconvolution analysis comparing estimated immune cell-type proportions between post-treatment LN and CT regions. D) Heatmap of representative lymphoid and T-cell-associated transcripts demonstrating compartment-specific transcriptional organization across LN and CT tissues.

### Digital spatial profiling identifies response-associated macrophage and B cell profiles

Multiplex immunofluorescence imaging using the Lunaphore COMET sequential immunofluorescence (seqIF) platform was performed to further resolve the cellular architecture underlying the transcriptional programs identified by GeoMx analysis. Imaging data was subsequently analyzed using the python-based SPACEc analysis pipeline for multiplexed image processing and analysis^36^. Representative H&E and COMET seqIF images demonstrated substantial spatial heterogeneity across response groups, including marked differences in immune infiltration and melanoma burden (Figure 3A). Melanoma-associated regions identified by HMB45 and SOX10 staining were more prominent in PR and NR tissues, whereas CR tissues demonstrated comparatively increased immune infiltration. Immune populations were defined using semi-supervised clustering per the SPACEc image analysis pipeline, and dimensionality reduction analyses identified distinct clusters of major immune and stromal populations (Figure 3B). Sub-clustering confirmed macrophage and T cell heterogeneity was present across all tissue cores, and the proportions of macrophage and lymphocyte populations differed across CR, PR and NR tissues (Figure 3C). Where NR and PR tissues had comparatively higher proportions of B cells and stroma compared to CR tissues. In contrast, CR tissues had higher proportions of CD8^+^ T cells, myeloid and intermediate as well as M2-like TAM populations.

**Figure 3.**
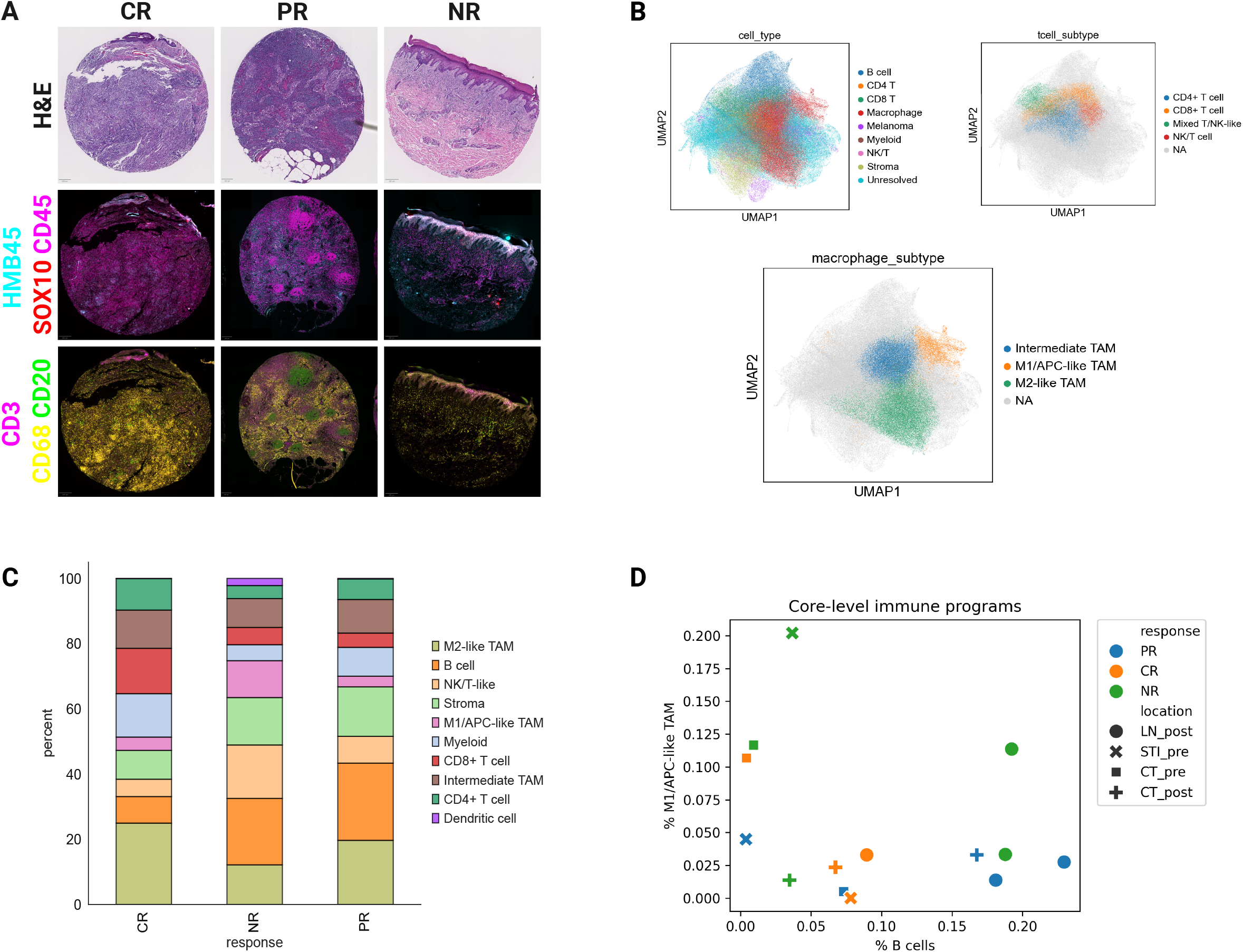
Digital spatial profiling identifies response-associated inflammatory macrophage populations. A) Representative H&E and multiplex immunofluorescence images from complete responder (CR), partial responder (PR), and non-responder (NR) tissues demonstrating spatial heterogeneity in immune infiltration and melanoma burden. B) UMAP visualization of SPACEc-derived single-cell clustering demonstrating major immune, stromal, and melanoma-associated cell populations across analyzed tissues. C) Relative abundance of annotated immune populations across CR, PR, and NR tissues, highlighting enrichment of inflammatory macrophage populations in responding tissues. D) Core-level comparison of M1/APC-like tumor-associated macrophage (TAM) and B-cell proportions across analyzed tissues.

To further examine the relationship between M1/APC-like TAM and B cell populations, tissue core-level immune programs were visualized according to treatment status, pathological response, and tissue region (Figure 3D). NR patient tissues showed clear separation in the proportions of M1/APC-like TAM to B cell populations based on therapy status; where higher proportions of M1/APC-like TAMs compared to B cell populations were seen in pre-T-VEC tissues, and the inverse was seen in post-T-VEC tissues. PR tissues showed low proportions of M1/APC-like TAMs and B cell pre-T-VEC, but higher proportions of B cells to M1/APC-like TAMs post-T-VEC. Whereas pre-T-VEC CR tissues had higher proportions of M1/APC-like TAMs to B cells, yet post-T-VEC CR tissues showed relatively balanced proportion of M1/APC-like TAM and B cell populations.

### HSV preferentially localizes with PD-1-high M1 TAMs enriched in CR tissues

Given the differences in proportions of M1/APC-like TAMs relative to B cell populations across CR, PR and NR tissues, downstream analyses also investigated the spatial relationship between HSV localization, macrophage phenotype and immune checkpoint status post-T-VEC therapy. To first confirm whether residual viral signal could be detected in post-T-VEC tissues, three HSV-associated antibodies were included in the COMET seqIF antibody panel to target HSV-associated proteins corresponding to distinct stages of the HSV lifecycle. These proteins included immediate-early (IE) protein ICP0, early protein ICP8, and the late glycoprotein HSVgD. Co-localization of all three HSV proteins was observed across post-T-VEC tissue cores (Figure 4A), and co-expression of ICP0, ICP8 and HSVgD was used to define HSV-positive cell states. HSV-positive (HSV^+^) cell status was overlayed onto previously annotated tumor and immune cell populations across CR, NR and PR tissues. Representative spatial maps highlighting the HSV expression relative to immune cell distributions are shown in Figure 4B.

**Figure 4.**
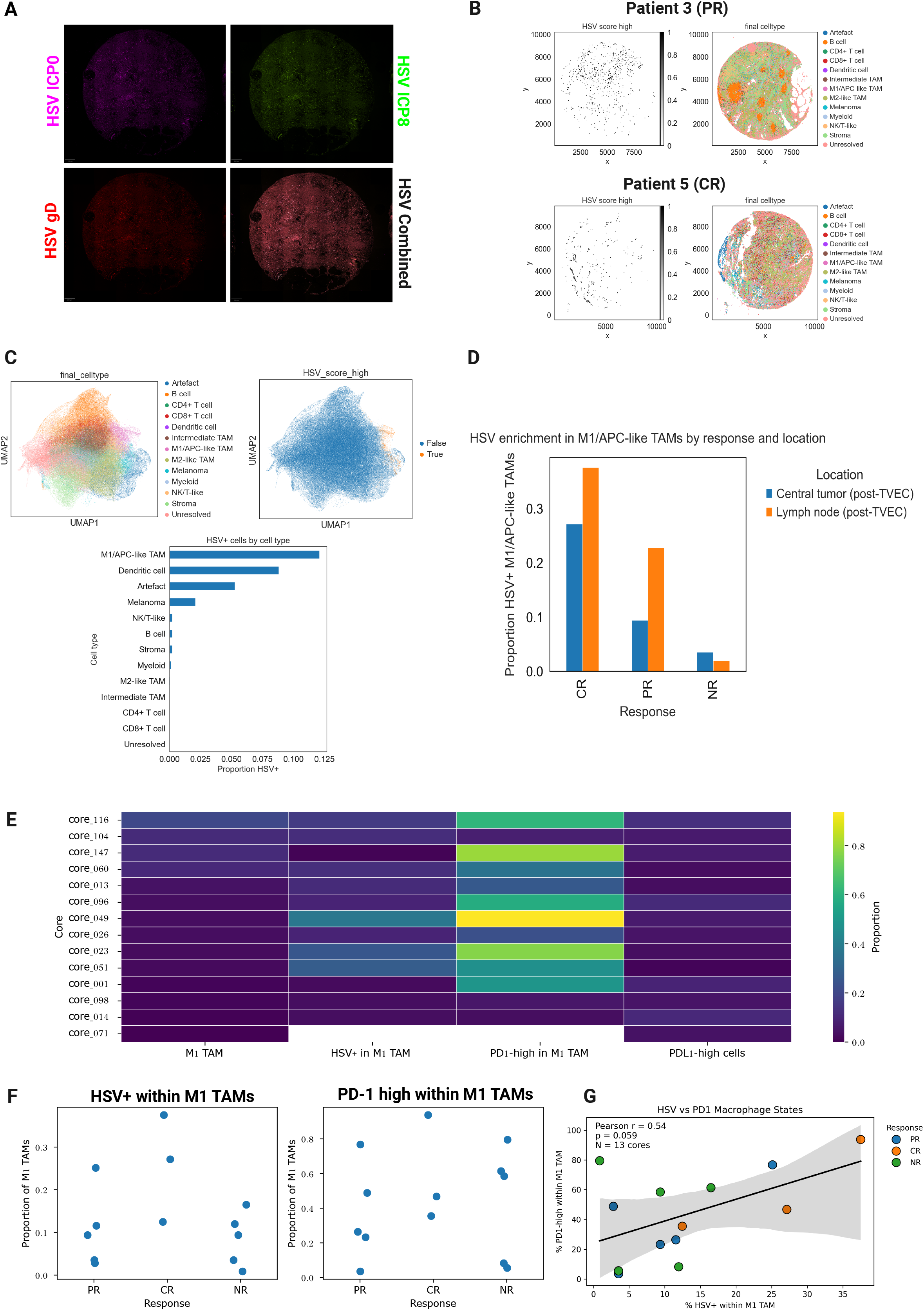
HSV preferentially localizes within PD-1-high inflammatory macrophages enriched in responding tissues. A) Representative multiplex immunofluorescence images demonstrating co-localization of three HSV-associated proteins corresponding to distinct stages of the viral lifecycle, including ICP0, ICP8, and HSVgD. Combined HSV-associated protein expression was used to generate a composite HSV score for downstream analyses. B) Representative spatial mapping of HSV-associated cells across PR and CR tissues following integration with SPACEc-derived single-cell annotations. C) UMAP visualization and quantification of HSV-associated cellular populations demonstrating preferential enrichment of HSV signal within M1/APC-like tumor-associated macrophages (TAMs). D) Relative abundance of HSV-positive M1/APC-like TAMs across pathological response groups and tissue compartments. E) Relative proportions of PD-1-high and PD-L1-high cellular populations across annotated immune cell types. F) Comparison of HSV-positive and PD-1-high M1/APC-like TAM populations across CR, PR, and NR tissues. G) Correlation between HSV-positive M1/APC-like TAM proportions and PD-1-high M1/APC-like TAM proportions across analyzed tissues.

Dimensionality reduction demonstrated that HSV^+^ cells clustered predominantly with macrophage-rich regions, and quantification of HSV distribution across annotated cell types confirmed M1/APC-like TAMs showed the highest proportion of HSV^+^ cells, followed by dendritic and melanoma cells (Figure 4C). Interestingly, the relative abundance of HSV^+^ M1/APC-like TAMs was enriched in CR compared to PR and NR tissues and were more abundant within LN tissues relative to post-T-VEC CT tissues across all three clinical response categories (Figure 4D). To investigate if increased HSV^+^ M1/APC-like TAM populations were also associated with increased immune checkpoint protein status, PD-1 and PD-L1 expression was also overlayed onto immune cell populations present across all tissue cores. Cell types were annotated as PD-1-high or PD-L1-high using a percentile fluorescence intensity threshold, and M1/APC-like TAMs represented one of the dominant PD-1-high immune populations across analyzed tissues, whereas comparatively lower PD-L1-high cells were observed within these macrophage populations (Figure 4E). The relative proportions of HSV^+^ M1/APC-like TAMs and PD-1-high M1/APC-like TAMs were compared across response groups, where CR tissues demonstrated enrichment of both HSV^+^ and PD-1-high M1 TAM populations relative to PR and NR tissues (Figure 4F). Correlation analyses further demonstrated a positive association between the proportion of HSV^+^ M1/APC-like TAMs and PD-1-high M1/APC-like TAMs across analyzed tissues, and was primarily driven by CR-associated regions, although this relationship nearly reached statistical significance (r = 0.59, p=0.059) (Figure 4G).

### Functional PD-1/PD-L1 interactive states and immune neighborhood organization differ across pathological responses

Functional oncology mapping (FuncOmap) analysis was also performed on tissue cores to determine if any differences could be observed in the level of functional PD-1/PD-L1 interactions post-T-VEC treatment across PR, NR and CR patients. FuncOmap incorporates fluorescence lifetime imaging microscopy (FLIM) and quantifies functional PD-1/PD-L1 interactive states occurring at a spatial resolution of 1-10nm. PD-1/PD-L1 interactive states are quantified as efficiency of energy transfer (Ef) which ranges from 0-50%, where the higher the Ef% the closer and stronger interaction between PD-1/PD-L1 and Ef <4% is considered non-interactive^38,39^. FuncOmap was performed on post-T-VEC CT and LN tissues from patients 3 (PR),4 (NR) and 5 (CR) which revealed that PD-1/PD-L1 interactive states were considerably higher in CT compared to LN tissues across all three patients, with upper quartile Ef (Q3) values ranging from 12-15% in CT regions compared to >4% Q3 Ef in LN regions (Figure 5A).

**Figure 5.**
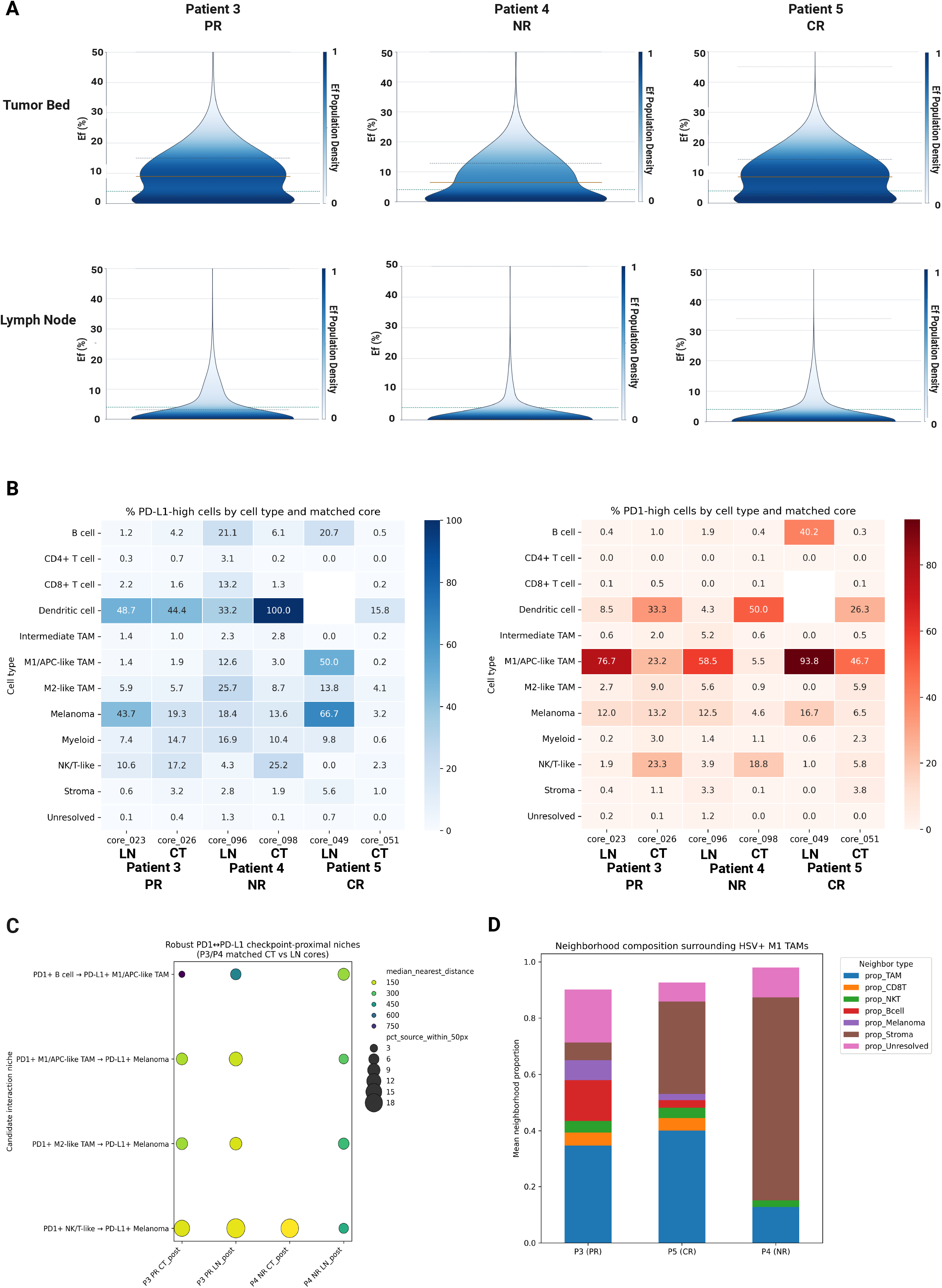
Functional PD-1/PD-L1 interaction states and immune neighborhood organization differ across pathological response groups. A) Violin plots showing global distribution of PD-1/PD-L1 interactive states (Ef%) quantified using FuncOmap analysis across matched post-treatment central tumor (CT) and lymph node (LN) tissues from patient 3 (n=27,000 data points), patient 4 (n=12,000 datapoints) and patient 5 (n=12,000 datapoints) as representatives of PR, NR, and CR responses respectively. B) Heatmaps showing the percentage of PD-1-high and PD-L1 high cells across annotated cell populations in matched post-TVEC central tumor bed (CT) and lymph node (LN) tissue cores collected from patients 3,4 and 5. Values represent the proportion of cells within each annotated lineage classified as PD-1-high or PD-L1-high based on per-core thresholding. C) Bubble plot summarizing the top PD-1/PD-L1 proximal ‘interactions’ identified through nearest-neighbor spatial screening in matched CT and LN tissues from patients 3 (PR) and 4 (NR). Bubble size indicates the percentage of PD-1-high source cells located within 50 pixels (px) of the indicated PD-L1-high target population, while color denotes median nearest-neighbor distance in pixels. Selected interactions represent biologically relevant PD-1/PD-L1 niches involving melanoma, TAM, NK/T-like, and B cell populations. D) Neighborhood composition analysis centered on HSV+ M1/APC-like TAM anchor cells in post-treatment CT cores. Stacked bar plots depict the normalized relative abundance of neighboring populations located within a 50-pixel radius of each anchor cell. Neighborhood compositions highlight distinct local and immune and tumor microenvironment across different pathological response groups.

Spatial proximity and cell neighborhood analyses were subsequently performed to investigate how tumor and immune cell niches of PD-1/PD-L1 expression may differ across matched post-T-VEC CT and LN tissues. The proportion of PD-1-high and PD-L1-high tumor and immune quantifications identified that M1/APC-like TAMs were predominantly associated with PD-1-high status, while PD-L1-high states were associated with melanoma and dendritic cell compartments (Figure 5B). Nearest-neighbor analysis identified the top recurring sources of potential PD-1/PD-L1 interaction patterns involving melanoma, TAM, NK/T and B cell populations in NR and PR tissues (Figure 5C). For CR tissues, anchored-based neighborhood analysis was performed centering on HSV^+^ M1/APC-like TAM populations. This revealed distinct immune and tumor neighborhood compositions surrounding HSV^+^ M1/APC-like TAMs in CR compared to NR and PR tissues (Figure 5D).

## Discussion

The goal of this study was to use integrated spatial transcriptomic and quantitative proteomic analyses to characterize immune remodeling and functional checkpoint interaction within the melanoma tumor immune microenvironment (TiME) following immunotherapy with oncolytic virus strain T-VEC. By applying this integrated approach to patients with known pathologic responses to T-VEC a secondary aim of the study was to identify if differences in immune spatial organization and functional checkpoint interactions could be associated to specific pathologic responses. While T-VEC induced widespread immune activation across tumor and LN tissues, quantitative spatial proteomic analyses demonstrated that pathologic response was not only associated with presence of certain immune populations, but also with the spatial organization of these populations relative to viral proteins in post-T-VEC tissue.

Initial transcriptomic analyses identified broad activation of lymphocyte- and myeloid-associated transcriptional programs in post-T-VEC tumor and LN tissues, consistent with systemic activation of anti-tumor immunity post oncolytic virotherapy. Successful and durable anti-tumor immune responses to immunotherapy have long been attributed to enhanced tumor infiltration and activity of CD8^+^ cytotoxic T lymphocytes (CTLs)^40,41^. However, increased activation of T lymphocyte-associated transcriptional programs post-T-VEC was found in LN but not tumor tissues and was accompanied by significant increases in CD4^+^ T cells. Although these T-VEC-induced changes in LN-based T cell transcription and infiltration were heterogenous across patient ROIs and pathologic responses. For post-T-VEC tumor tissues, the most prominent transcriptional changes observed were increased immunoglobulin (Ig)-associated genes, whose expression was consistently elevated in patients with partial pathologic response (PR). This Ig signal was not accompanied by a coordinated enrichment of canonical B cell transcriptional programs or significant difference in B or plasma cell proportions. Instead, immunoglobulin-associated transcription showed only moderated association with B-lineage abundance and was heterogeneous across individual patients and ROIs. Collectively, these results highlighted region-specific immune remodeling post-T-VEC across tumor and LN tissues potentially arising from localized B and T lymphocyte niches that could not be fully resolved using bulk transcriptions and cellular deconvolution.

COMET seqIF analyses were subsequently performed across whole tumor and LN tissue cores to validate and spatially contextualize post-T-VEC immune remodeling initially identified within GeoMx DSP ROIs. Substantial B cell, macrophage and T cell heterogeneity was observed across patient pathologic response categories, where the relative abundance of macrophages compared to B cells was higher for CR compared to PR and NR patients. These results are consistent with prior T-VEC studies in melanoma and cutaneous basal cell carcinoma (BCC), where T-VEC-induced remodeling included increased infiltration of macrophage, T cell and B cell populations. For macrophage and T cell populations, increased M1 tumor associated macrophages (TAMs) and CD8^+^ T cells post-T-VEC were associated with CR, however the relationship between B cells and PR/NR are less understood^42,43^. In BCC, T-VEC induced an expansion of B cell and plasma cell subsets which was heterogeneous between patients and was not significant between response groups ^42^. However, single cell RNA and B cell receptor sequencing identified that IGHG1^+^ plasma cell clones were hyper-expanded and associated with CR.

Multiplex immunofluorescence (mIF) also confirmed this population of IgG1^+^CD138^+^ plasma cells was enriched in CR patients, but it is unclear if mIF was able to detect any difference in B cell or Ig-heavy B cell populations in this study. Conversely, Mulder et al.’s study investigating humoral responses to T-VEC in 30 patients with stage III-IV metastatic melanomas found post-T-VEC CD138^+^ plasma cell populations increased relative to CD8^+^ T cells, however, no relation to durable response was observed. A relationship between B cell populations and ICB-resistance has also been reported, where IGHA1 and IGHG1 memory B cell proportions were increased relative to plasma cells in ICB-resistant melanomas^43^, however further studies are also needed to determine the role of these B memory cell functions in driving ICB resistance. In the context of T-VEC therapy, our future studies will also incorporate additional plasma cell markers to investigate if and how the distribution of B cell verses plasma cells may differ according to pathologic response and tissues region to further explore how differences in the relative abundance of these two cell populations may affect PD-1/PD-L1 interaction states quantified by FuncOmap analysis.

Another crucial component of understanding T-VEC anti-tumor immunity is determining how immune populations are spatially organized relative to HSV-associated regions within the melanoma TiME. Yet the majority studies using spatial omics to characterize T-VEC-induced immune remodeling do not incorporate anti-HSV antibodies or DNA probes. One exception is Mulder et al.’s study, where the presence of HSV-1 protein and DNA post-T-VEC was analyzed using conventional immunohistochemistry (IHC) and real-time PCR^43^. While IHC was unable to detect HSV across any post-T-VEC patient samples, PCR detected very low amounts of HSV-1 DNA, leading the authors to conclude this was evidence of cleared HSV-1 infection. However, as no HSV-1 PCR or IHC data is shown, it is difficult to determine if infections were equally cleared across all patient samples or differed according to response status. The limited levels of HSV detection by IHC may be due to IHC analysis being performed using antibodies only targeting intermediate early (IE) HSV protein ICP8, whose expression traditionally peaks within 6-12 hours post infection *in vitro*^44^ and less is known of the stability of IE protein dynamics in FFPE preserved human tissues.

Multiple early, IE and late anti-HSV antibodies were used in this study to maximize detection of HSV across post-T-VEC tissues, which identified HSV localization was most strongly enriched within M1/APC-like TAM populations relative to T and B lymphocyte populations. The enrichment of these HSV^+^ M1/APC-like TAMs in CR compared to PR/NR tissues further supports a role for M1 macrophages as key mediators of OV-associated anti-tumor immune responses. While macrophages are well established as central mediators of anti-viral immunity to wild-type HSV infection^45^, the role of M1 TAMs in shaping immune responses to engineered oncolytic HSVs in melanomas has been less extensively characterized. Particularly regarding their role in driving systemic immune responses in sentinel LN (SLN), as the benefit of SLN biopsies in the context of neoadjuvant immunotherapy is still a source of heavy clinical debate^46^. Our finding that M1/APC-like TAMs were a major source of PD-1 expression and were enriched in LN compared to tumor tissues in CR/PR tissues suggests that M1 TAM populations could be key regulators of systemic anti-tumor responses to T-VEC and may also provide localized niches for checkpoint interactions within responding tissues.

Integration of FuncOmap PD-1/PD-L1 checkpoint interaction analysis with COMET seqIF analysis highlighted that while PD-1/PD-L1 interaction states were consistently elevated in post-T-VEC tumors compared to LN tissues across all response groups, the spatial organization of PD-1/PD-L1-associated immune cells differed across CR, NR and PR tissues. Where neighborhoods surrounding HSV^+^ M1 TAM PD-1-high niches in CR tissues were dominated by other TAM cell populations. In contrast, PR and NR tissues retained melanoma-centered PD-1/PD-L1 associated immune niches enriched with B cell and M2-like TAM populations. Altogether, these findings indicate that pathological response to T-VEC therapy may depend less on only checkpoint protein interactions but requires their correlation with spatial and cellular context in which these interactions occur.

Several limitations should be considered when interpreting these findings. First, this study cohort was relatively small and included noticeable inter- and intra-patient heterogeneity, limiting the ability to draw definitive conclusions regarding the role of specific immune populations across response groups. In addition, transcriptomic cellular deconvolution is inherently dependent on the cell state reference matrix used and may incomplete capture transitional or functionally activated immune populations within T-VEC-treated tissues. Despite these limitations, the integration of transcriptomics, seqIF and FuncOmap enabled spatial and functional characterization of immune niches not easily identifiable by one modality alone. Future studies incorporating larger cohorts, inclusion of additional B and plasma-cell specific probes and FuncOmap region-specific seqIF profiling will be important to validate these findings and determining whether HSV-associated M1 TAM and B cell niches may represent biomarkers or therapeutic targets for OV therapy. Overall, these findings demonstrate the value of integrated spatial and quantitative functional immune profiling for identifying mechanisms of response and resistance to OV therapy within melanoma TiMEs.

## Conclusion

Integrated spatial transcriptomic and functional proteomic analyses demonstrated that T-VEC therapy is associated with substantial remodeling of the melanoma tumor bed and lymph node microenvironments. While adaptive immune and B cell-associated transcriptional programs were broadly induced following T-VEC, higher-resolution spatial analysis identified enrichment of HSV associated inflammatory macrophage populations exhibiting immune checkpoint PD-1/PD-L1-associated phenotypes in responding tissues. FuncOmap and spatial neighborhood analyses further demonstrated that organization of PD-1/PD-L1 interactive states and expression differed according to pathological response, with NR and PR tissues retaining melanoma-centered interaction landscapes with B cell and M2-TAM-driven niches and CR tissues exhibiting M1/APC-like TAM centered neighborhoods following tumor clearance. Together, these findings highlight distinctive immune profiles for mediating response and resistance to oncolytic virotherapy and highlight the importance of integrated spatial multi-omic approaches for resolving quantitative functional immune architectures within melanoma TiMEs.

## Acknowledgements

This work was generously supported by the John and Marva Warnock Endowed Scholar Fund, as well as by the Stanford Cancer Institute’s Institutional Research Grant from the American Cancer Society, Grant # IRG-19-218-01.

